# Antigenic variation by switching inter-chromosomal interactions with an RNA splicing locus in trypanosomes

**DOI:** 10.1101/2020.01.27.921452

**Authors:** Joana Faria, Vanessa Luzak, Laura S.M. Müller, Benedikt G. Brink, Sebastian Hutchinson, Lucy Glover, David Horn, T. Nicolai Siegel

## Abstract

Highly selective gene expression is a key requirement for antigenic variation in several pathogens, allowing evasion of host immune responses and maintenance of persistent infections. African trypanosomes, parasites that cause lethal diseases in humans and livestock, employ an antigenic variation mechanism that involves monogenic antigen expression from a pool of >2500 antigen coding genes. In other eukaryotes, the expression of individual genes can be enhanced by mechanisms involving the juxtaposition of otherwise distal chromosomal loci in the three-dimensional nuclear space. However, trypanosomes lack classical enhancer sequences or regulated transcription initiation and the monogenic expression mechanism has remained enigmatic. Here, we show that the single expressed antigen coding gene displays a specific inter-chromosomal interaction with a major mRNA splicing locus. Chromosome conformation capture (Hi-C), revealed a dynamic reconfiguration of this inter-chromosomal interaction upon activation of another antigen. Super-resolution microscopy showed the interaction to be heritable and splicing dependent. We find that the two genomic loci are connected by the antigen exclusion complex, whereby VEX1 associated with the splicing locus and VEX2 with the antigen coding locus. Following VEX2 depletion, loss of monogenic antigen expression was accompanied by increased interactions between previously silent antigen genes and the splicing locus. Our results reveal a novel mechanism to ensure monogenic expression, requiring the spatial integration of antigen transcription and mRNA splicing in a dedicated compartment. These findings suggest a new means of post-transcriptional gene regulation.

## Main Text

Monogenic expression, the expression of a single gene from a large gene family, is essential for several important biological processes. One of the most striking examples of such regulation is the expression of a single odorant receptor from more than 1400 genes in mammalian olfactory sensory neurons ^1^. Likewise, monogenic expression is a key feature of antigenic variation, an immune evasion strategy used by pathogens such as *Plasmodium falciparum* or *Trypanosoma brucei*. Antigenic variation refers to the capacity of an infecting organism to systematically alter the identity of proteins displayed to the host immune system^2^. How pathogens ensure the exclusive expression of only one antigen from a large pool of antigen coding genes remains one of the most intriguing questions in infection biology.

In *T. brucei*, a unicellular parasite responsible for lethal and debilitating diseases in humans and animals, 10 million copies of a single variant surface glycoprotein (VSG) isoform are exposed on the surface of the parasite. The exclusive expression of only one VSG gene per cell and the periodic switching of the expressed VSG gene allow the parasite to evade the host immune system and to maintain persistent infections ^3,4^. While the *T. brucei* genome encodes for >2600 VSG isoforms, in the bloodstream of the mammalian host, a VSG gene can only be transcribed when located in one of ∼15 VSG expression sites. Those bloodstream expression sites are polycistronic transcription units located adjacent to telomeres on different chromosomes. Each bloodstream expression site contains an RNA polymerase I (Pol I) promoter, followed by several expression site associated genes (ESAGs) and a single VSG gene ^5^.

Notably, Pol I transcription initiates at all VSG expression site promoters, but transcription elongation and transcript processing are highly selective and limited to just one expression site at a time ^6,7^. As a result, the single active VSG gene is expressed as the most abundant mRNA and protein in the cell; 5-10% of the total in each case. Why transcription is aborted at all but one expression site is not known, but it has been shown that only transcripts from the actively transcribed expression site are processed into mature mRNA. In trypanosomes, mRNA maturation involves *trans*-splicing, a process that adds a common spliced leader sequence to each pre-mRNA and is coupled to polyadenylation ^8^. In addition, the proximity of individual genes to nuclear condensates composed of splicing factors has recently been proposed to play a role in gene expression regulation in mammals ^9,10^. Thus, regulated access to RNA maturation compartments may represent an evolutionary conserved strategy for gene expression control.

One mechanism to ensure monogenic expression is the juxtaposition of otherwise distal chromosomal loci in the three-dimensional nuclear space. In particular, specific interactions between promoter and enhancer sequence elements can ensure the selective regulation of individual genes. Although classic enhancer structures appear to be absent in many unicellular eukaryotes such as trypanosomes, several observations suggest that a specific genome organization is required for monogenic VSG expression. The single active VSG gene is transcribed in an extranucleolar Pol I compartment known as the expression site body ^11^. In those very rare cases (<10^-8^) where two VSG genes are simultaneously active, both co-localize at the expression site body ^12,13^. In addition, the transcribed chromosome core regions and the sub-telomeric regions coding for the large reservoir of silent VSG genes, appear to fold into structurally distinct compartments ^5^, similar to active A and silent B compartments described in mammalian cells ^14^. While the nature of the expression site body has remained enigmatic, a protein complex specifically associated with the active VSG gene was identified recently. VSG-exclusion 1 (VEX1) emerged from a genetic screen for allelic exclusion regulators ^15^ while VEX2 was affinity-purified in association with VEX1 ^16^. The bipartite VEX protein complex maintains mutually exclusive VSG expression ^16^ but it remains unclear how these proteins exert their function. In this study we aimed to identify the mechanism that connects RNA maturation, genome architecture and the VEX complex to ensure monogenic antigen expression.

Given the well-characterized role of promoter-enhancer interactions in the selective regulation of genes, we set out to identify specific DNA-DNA interactions with a regulatory role in monogenic VSG expression. To this end we used a *T. brucei* culture homogenously expressing a single VSG gene for chromosome conformation capture (Hi-C) analysis. In addition, we employed the mHi-C analysis pipeline, which allowed us to retain many multi-mapping reads and greatly increased the read coverage across repetitive regions of the genome ^17^.

In order to visualize specific interaction patterns of loci of interest (viewpoints) in the Hi-C dataset, we applied a virtual 4C analysis pipeline to extract genome-wide interaction profiles for chosen viewpoints. To identify VSG gene specific interaction patterns, we chose the active and several inactive VSG genes located in expression sites as viewpoints and plotted the extracted virtual 4C interaction data onto the genome. As expected, we observed a distance-dependent decay of intra-chromosomal interactions between each viewpoint and its upstream and downstream genomic region (**Fig. 1a** **and Extended Data Fig. 1a**).

**Fig. 1:**
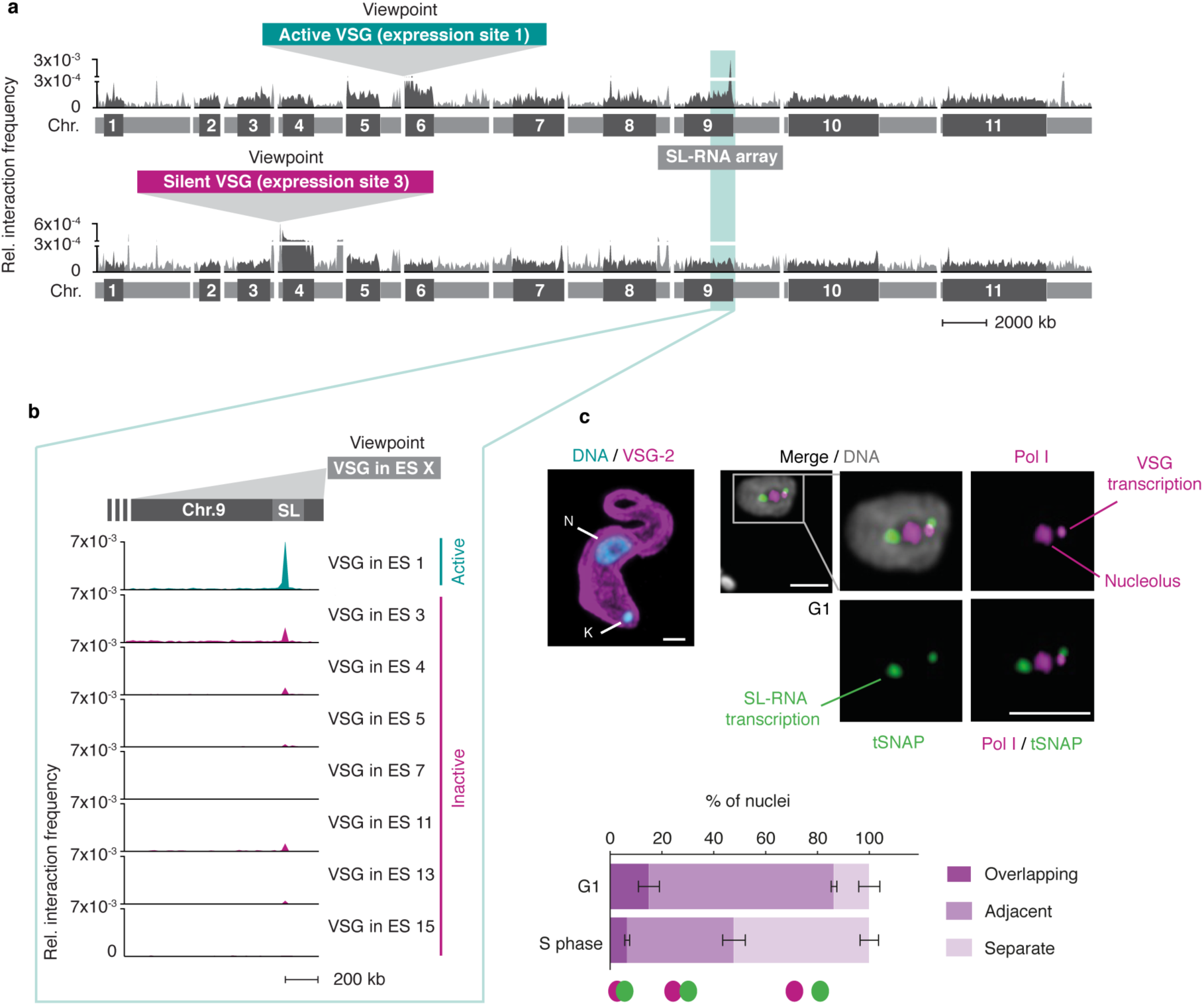
The active VSG expression site (ES) stably interacts with the spliced leader RNA (SL) array. **a,** Hi-C (virtual 4C) analysis, viewpoints: active VSG gene in ES1 (*VSG-2*, top panel) and silent VSG gene in ES 3 (*VSG-6*, bottom panel). Relative interaction frequencies between the viewpoint and the 11 megabase chromosomes are shown. Chromosome cores, dark grey; sub-telomeric regions, light grey. The hemizygous sub-telomeric regions are displayed in the following order: 5’(haplotype A)–5’(haplotype B)–diploid chromosome core–3’(haplotype A)– 3’(haplotype B). Bin size 50 kb. **b,** Virtual 4C analysis, viewpoints: active VSG gene in ES 1 and inactive VSG genes in ES 3, 4, 5, 7, 11, 13 and 15. Relative interaction frequencies between the viewpoint and the SL-RNA locus on the right arm of chr. 9 is plotted. Bin size 20 kb. The analyses in **a**-**b** are based on Hi-C experiments with *VSG-2* expressing cells (n=2, average interaction frequencies are shown). **c,** Immunofluorescence-based colocalisation studies of tSNAP^myc^ (SL-RNA locus marker – SL-RNA transcription compartment) and a nucleolar and active-*VSG* transcription compartment marker (Pol I, largest subunit) using super resolution microscopy. The stacked bar graph depicts proportions of G1 or S phase nuclei with overlapping, adjacent or separate signals for the SL-RNA and VSG transcription compartments. Values are averages of three independent experiments and representative of two independent biological replicates (≥100 G1 or S phase nuclei); error bars, SD. Detailed *n* and *p* values are provided in Data S1 sheet 3. DNA was counter-stained with DAPI; the images correspond to maximal 3D projections of stacks of 0.1 μm slices; scale bars 2 μm. N, nucleus; K, kinetoplast (mitochondrial genome).

Strikingly, we found the active *VSG-2* gene located on chr. 6 in expression site 1 to very frequently interact with a single, distinct locus on chr. 9 (**Fig. 1a-b**). Levels of interaction frequency were higher than intra-chromosomal interactions of *VSG-2* with its genomic location on chr. 6, pointing to a strong and stable inter-chromosomal interaction. The locus on chr. 9 interacting with the active VSG gene is the SL-RNA array, a genomic locus essential for RNA maturation. This locus contains a cluster of ∼150–200 tandemly repeated genes encoding the spliced leader RNA (SL-RNA). SL-RNA is an RNA Pol II-transcribed ncRNA that is *trans*-spliced to the 5’-end of all trypanosome mRNAs, conferring the 5’-cap structure required for RNA maturation, export and translation ^8^. Conversely, VSG genes residing in inactive expression sites interacted less frequently or at background levels with the SL-RNA locus (**Fig. 1a-b** **and Extended Data Fig. 1a**). In agreement with these observations, when we chose the SL-RNA locus as viewpoint, we found it to interact more frequently with the active VSG expression site than with any inactive VSG expression site (**Extended Data Fig. 1b**). Thus, the Hi-C analysis revealed a strong and selective interaction between the Pol I-transcribed active VSG gene and the Pol II-transcribed SL-RNA locus located on a different chromosome.

To visualize the spatial proximity between the active VSG gene and the SL-RNA locus at the level of individual cells and with an independent assay, we performed super-resolution immunofluorescence microscopy. The site of VSG transcription is characterized by an extra-nucleolar accumulation of RNA Pol I ^11^. The site of Pol II-transcribed SL-RNA is marked by an accumulation of the small nuclear RNA-activating protein complex (tSNAPc), an RNA Pol II promoter-binding transcription factor ^18^. Both the VSG transcription and the SL-RNA transcription compartments appear to have diameters of approx. 300 nm, within a *T. brucei* nucleus with a diameter of approx. 2 μm. Super-resolution microscopy revealed one VSG transcription compartment, reflecting the hemizygous active sub-telomeric *VSG-2*, and two separate SL-RNA transcription compartments, reflecting the diploid SL-RNA arrays located in the core of chr. 9 (**Fig. 1c**). By scoring nuclei for overlapping, adjacent or separate VSG and SL-RNA transcription compartments, we found that one of the SL-RNA transcription compartments was adjacent to the VSG transcription compartment in the majority of cells (**Fig. 1c**). However, during DNA replication in S phase, the VSG and SL-RNA transcription compartments were detected in separate locations in >50% of nuclei (**Fig. 1c** **and Extended Data Fig. 1c**). Therefore, throughout this study, immunofluorescence assays (IFAs) were subsequently performed in G1 cells, unless indicated otherwise. Taken together, IFAs supported the findings made by Hi-C, suggesting that the Pol I VSG transcription compartment interacts with one of the Pol II SL-RNA transcription compartments. Further, they suggest that the interaction between both compartments is resolved during S phase and successfully re-established after replication.

To determine whether the interaction with the SL-RNA transcription compartment is specific for the active VSG gene, and therefore changes following a VSG switching event, we performed Hi-C experiments using an isogenic *T. brucei* cell line expressing a different VSG isoform, *VSG-13* (**Fig. 2a**) ^19^. *VSG-13* resides within expression site 17, which is located on one of five intermediate-sized chromosomes. The presence of co-transcribed resistance markers upstream of *VSG-2* in expression site 1 and *VSG-13* in expression site 17 allowed us to specifically select for parasites expressing *VSG-13* through drug selection (**Fig. 2a**). The exclusive activity of expression site 1 or 17 was verified by RNA-seq (**Fig. 2a**).

**Fig. 2:**
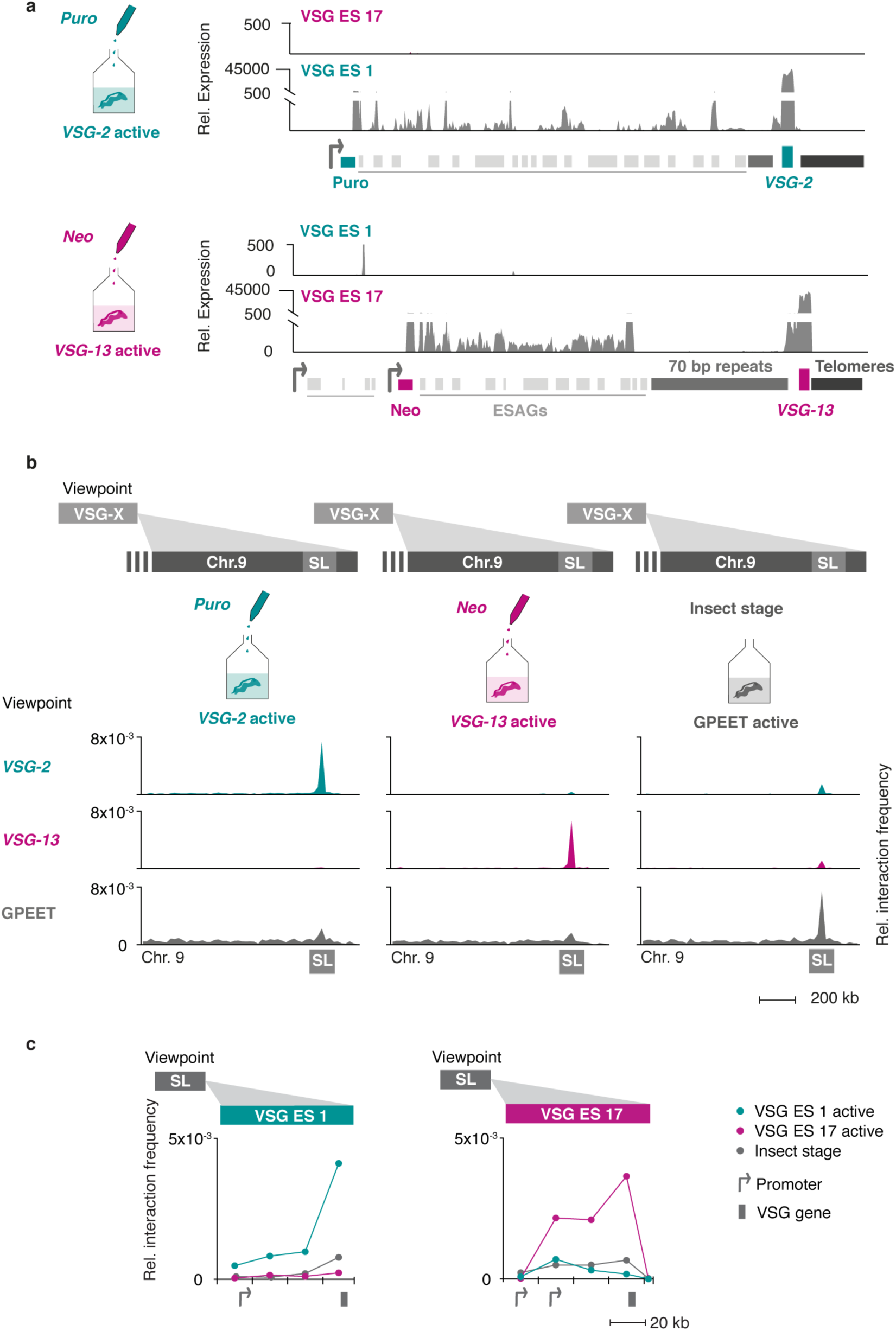
The interaction between the active VSG gene and the SL-RNA locus is dynamic and changes during a switch in VSG expression. **a,** Transcriptome analyses of isogenic cell lines after selection with puromycin (active: *VSG-2*), top panel, or neomycin (active: *VSG-13*), bottom panel. For each condition, the average of three biological replicates is plotted. **b,** Hi-C (virtual 4C) analysis, viewpoints: *VSG-2*, *VSG-13*, GPEET gene array (chr. 6). Relative interaction frequencies between the viewpoint and the SL-RNA locus on chr. 9 are plotted. Bin size 20 kb. **c,** Hi-C (virtual 4C) analysis, viewpoint: SL-RNA locus (chr. 9). Relative interaction frequencies between the viewpoint and VSG ES 1 or ES 17 are shown. Bin size 20 kb. The analyses in **b** and **c** are based on Hi-C experiments with cells expressing *VSG-2*, *VSG-13* or insect stage cells expressing procyclin genes (n=2, the average is shown).

Hi-C analysis revealed that *VSG-2 -* SL-RNA interactions dropped 20-fold to average inter-chromosomal interaction levels in parasites expressing *VSG-13*, while interactions between the newly activated *VSG-13* and the SL-RNA locus increased 36-fold (**Fig. 2b, left and middle panel**). We found that upon activation of each expression site, the bin harboring the respective active VSG gene displayed the strongest interaction with the SL-RNA locus, suggesting that the VSG gene itself, not its promoter, interacts with the splicing locus (**Fig. 2c**). In addition, we detected decreased interaction of the inactivated *VSG-2* gene with the transcribed chromosome cores, while the activated *VSG-13* gene displayed increased interaction with chromosome cores (**Extended Data Fig. 2a**). This observation indicates that activation of a VSG gene is intimately linked to a transition from a silent to an actively transcribed compartment within the nucleus. Conversely, VSG gene inactivation results in a transition from an active to an inactive nuclear compartment.

To further explore the relationship between SL-RNA interaction frequency and gene expression, we performed Hi-C analyses using insect stage parasites that do not express any VSGs, but instead express a different group of surface antigens called procyclin genes. Confirming the importance of the SL-RNA interaction, the GPEET and EP1 procyclin genes displayed increased interaction frequency with the SL-RNA locus upon activation in insect stage cells (**Fig. 2b, right panel, and Extended Data Fig. 2b**). Like VSG genes, procyclin genes are transcribed by RNA Pol I at high levels and require efficient *trans*-splicing for mRNA maturation. Thus, Hi-C analyses of *T. brucei* cell lines expressing different antigens indicate that interactions with the SL-RNA locus are dynamic and specific for the actively transcribed antigen coding genes.

Previously, we had shown that the bipartite VEX-complex is associated with the actively transcribed VSG gene and maintains monogenic VSG expression but that VEX1 and VEX2 only partially overlap each other ^16^. Given a similar juxtaposition of the VSG transcription and the SL-RNA transcription compartments, we sought to investigate the relationship between the VEX complex and these transcription compartments in more detail. Using optimized immunofluorescence staining protocols and super-resolution microscopy, we were able to detect two VEX1 foci in the majority of G1 cells (55 +/-4 %). These VEX1 signals specifically co-localized with the SL-RNA transcription compartments (**Fig. 3a**). In contrast, the majority of G1 cells (97 +/-1 %) only had one VEX2 focus, which specifically co-localized with the VSG transcription compartment (**Fig. 3b**). As expected, one VEX1 focus was adjacent to the VSG transcription compartment (**Extended Data Fig. 3a**) while the VEX2 focus was adjacent to one of the two SL-RNA transcription compartments (**Extended Data Fig. 3b**). Thus, our IFAs revealed association of VEX1 with the SL-RNA transcription compartments and VEX2 with the VSG transcription compartment. To verify a specific interaction between VEX1 and the SL-RNA locus, we reanalyzed published VEX1-ChIP-seq data, previously only mapped to VSG expression sites ^16^. The ChIP-seq data revealed a striking enrichment of VEX1 at the SL-RNA locus that was greater than at any other gene, including the active VSG gene (**Fig. 3c, Extended Data Fig. 3c; Data S1, sheet 1**). These data suggest that the VEX-complex connects the VSG and SL-RNA transcription compartments.

**Fig. 3:**
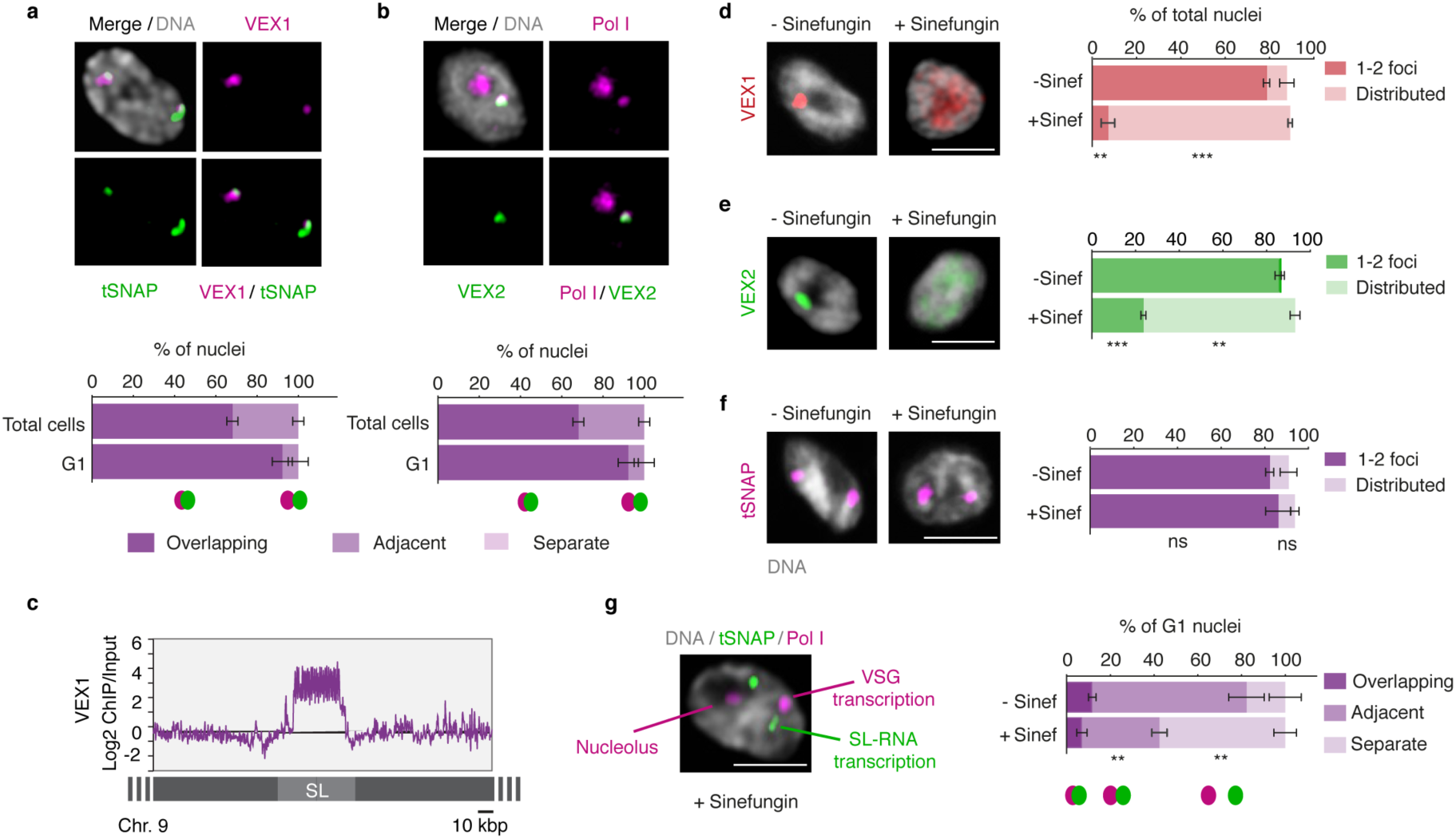
The VEX complex associates with both the active-*VSG* and the Spliced Leader (SL)-locus in a splicing-dependent manner. **a-b,** Immunofluorescence-based colocalisation studies of VEX1^myc^ / tSNAP^GFP^ and ^GFP^VEX2 / Pol I. tSNAP and Pol I are used as markers for the SL-RNA and VSG transcription compartments, respectively. The stacked bar graphs depict proportions of nuclei with overlapping, adjacent or separate signals: values are averages of four (**a**) or two (**b**) independent experiments (≥200 nuclei for total cell counts; ≥100 nuclei for G1 phase). **c,** VEX1^myc^ chromatin immunoprecipitation followed by next generation sequencing (ChIP-Seq) analysis. The graph depicts log2 fold change of ChIP signal versus input sample across the SL-RNA locus. Bin size 300 bp. **d-f,** Immunofluorescence analysis of VEX1^myc^ (**d**), ^myc^VEX2 (**e**) and tSNAP^myc^ (**f**) before and after sinefungin treatment (5 μg ml^-1^ for 30 min at 37°C). Cells displaying no detectable signal (<10%) were excluded. Values are averages of two independent experiments (≥200 nuclei each). **g,** Immunofluorescence-based colocalisation studies of the SL-RNA transcription (tSNAP^myc^) and the VSG transcription (Pol I, large subunit) compartments following treatment with sinefungin. The stacked bar graph depicts proportions of G1 nuclei with overlapping, adjacent or separate signals and values are averages of two independent experiments and two biological replicates (≥100 G1 nuclei). The studies in **a-b** / **d-g** were undertaken using super resolution microscopy and the images correspond to maximal 3D projections of stacks of 0.1 μm slices; DNA was counter-stained with DAPI; scale bars 2 μm. In **d-g,** a two-tailed paired Student’s *t*-test was used to compare non-treated versus treated nuclei for each category: ns, non-significant; **, *p* < 0.01; ***, *p* < 0.001. Experiments in **a-b** / **d-g** are representative of at least two independent biological replicates and detailed *n* and *p* values are provided in Data S1 sheet 3.

Although VSG and SL-RNA transcription compartments separate during S phase (**Fig. 1c**), VEX1 does not separate from the SL-RNA transcription compartment and VEX2 does not separate from the VSG transcription compartment (**Fig. 3a and b**). Also, consistent with the loss of VSG expression in insect-stage cells, SL-RNA transcription compartments can still be identified through tSNAP localisation, while the VEX and VSG transcription compartments are reorganized and lost, respectively (**Extended Data Fig. 3d-e**). These results indicate that VEX2 marks the VSG transcription compartment in a developmental stage specific manner, which in bloodstream stage cells may facilitate the re-establishment of compartment connectivity to propagate expression of a specific antigen. Consistent with this idea, VEX complex reassembly after DNA replication is dependent upon CAF-1 histone chaperone function ^16^.

Given the close spatial proximity between the site of VSG transcription and the site of SL-RNA transcription, we next questioned whether the splicing process itself impacts the connection between these compartments. We found that inhibition of *trans*-splicing with sinefungin ^20^ disrupted both VEX1 (**Fig. 3d**) and VEX2 (**Fig. 3e**) localization within 30 min, while the tSNAP transcription factor was not affected under the same conditions (**Fig. 3f**). Notably, inhibition of splicing by sinefungin also disrupted the connection between the VSG and SL-RNA transcription compartments, revealed by separation of the Pol I and tSNAP signals (**Fig. 3g**); neither the VEX, nor the tSNAP protein levels were affected by sinefungin treatment (**Extended Data Fig.3f**). Thus, VEX protein localization and the juxtaposition of the VSG and the SL-RNA transcription compartments are dependent on mRNA splicing activity.

Next, we aimed to investigate the mechanism by which the VEX complex ensures monogenic VSG expression. Previously, we found that VEX2 depletion leads to a strong activation of VSG genes located in previously silent expression sites ^16^. Thus, following our observation that the VEX-complex spans the VSG transcription and the SL-RNA transcription compartments, we sought to determine whether VEX2 functions as a connector or as an exclusion factor. That is, whether following VEX2 depletion the active VSG gene loses connectivity to the SL-RNA transcription compartment or whether previously inactive expression sites start to interact with the SL-RNA transcription compartment. To test these models, we analyzed the distance between the VSG transcription compartment and the SL-RNA transcription compartment following depletion of VEX-complex components (**Extended Data Fig. 4a-b**). IFA data revealed a disruption of compartment connectivity in 45% of G1 nuclei following 12 hours of VEX2 knockdown (P <0.001) and 60% following VEX1 - VEX2 double-knockdown (P <0.01) (**Fig. 4a, Extended Data Fig. 4c**); the mean sub-compartment distances were 60 nm in control cells and 167 nm following VEX-complex knockdown (**Fig. 4a-b, Extended Data Fig. 4d**). At later time points after the induced VEX2 knock down, the RNA Pol l signal disperses ^18^, indicating that VEX2 sustains a local reservoir of Pol I at the active VSG gene. Compartment connectivity was not significantly disrupted following VEX1-knockdown.

**Fig. 4:**
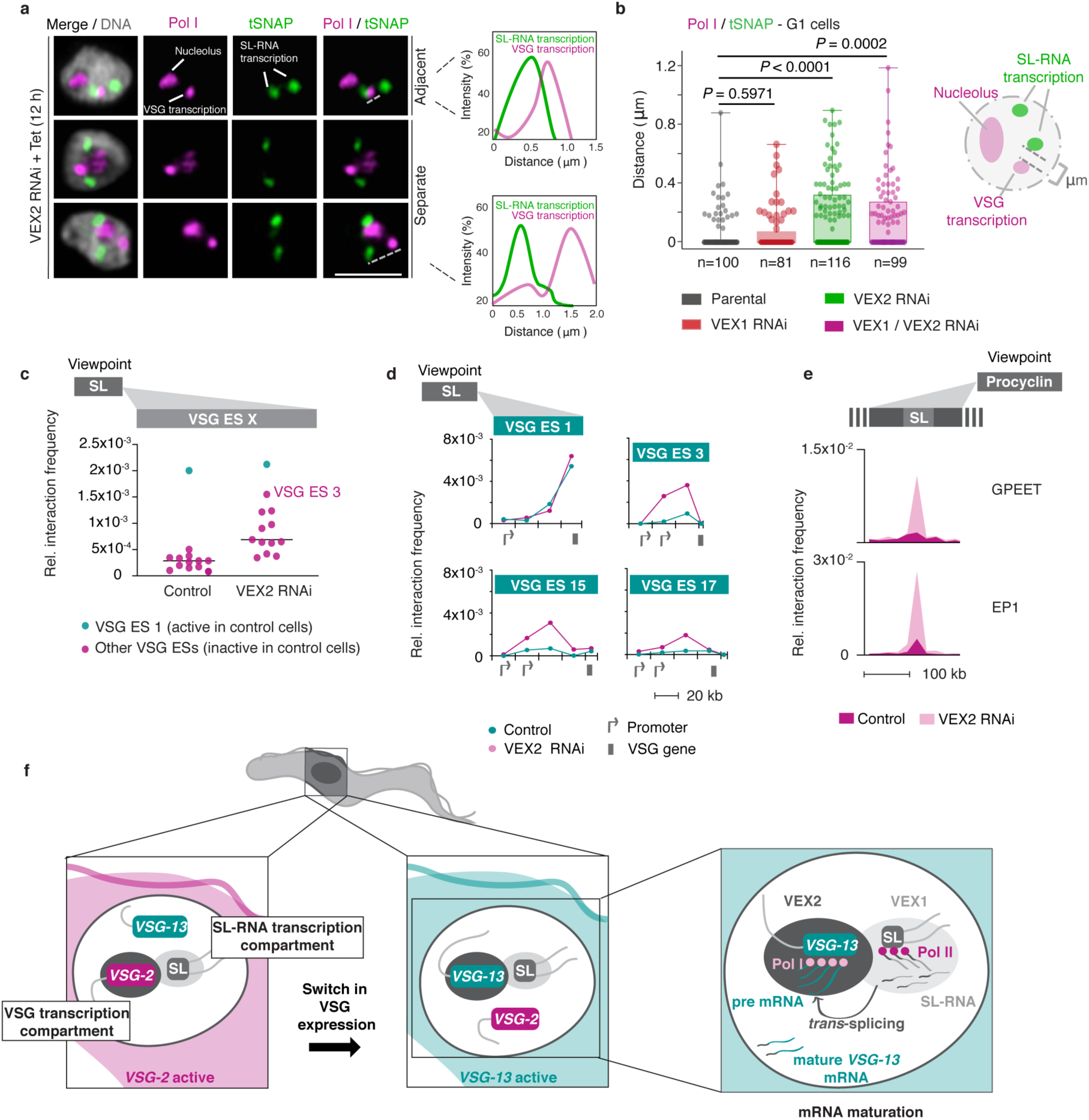
The exclusive association between the active VSG and the *SL*-locus is VEX2-dependent. **a-b,** Immunofluorescence and super resolution microscopy based colocalisation studies of tSNAP^myc^ (SL-RNA transcription compartment) and Pol I (nucleolus and active-VSG transcription compartment) following VEX1, VEX2 and VEX1/VEX2 knockdown. **a,** Representative images, all nuclei are G1. On the right-hand side, two representative histograms depict the distribution of signal intensity across the distance indicated by the dashed lines. Images correspond to maximal 3D projections of stacks of 0.1 μm slices; DNA was counter-stained with DAPI; scale bars 2 μm. **b,** The box plot depicts the distance between the active-VSG and the nearest SL-RNA transcription compartment (≥ 81 G1 nuclei) following VEX1, VEX2 and VEX1/VEX2 knockdown; the centre lines show the medians; box limits indicate the 25th and 75th percentiles; whiskers extend from maximal to the minimal values; all data points are shown. The distances for each knockdown condition were compared to the parental cell line using a two-tailed unpaired Student’s *t*-test. **c,** Hi-C (virtual 4C) analysis, viewpoint: SL-RNA locus (chr. 9). Relative interaction frequencies between the viewpoint and the VSG expression sites are shown before and after VEX2 knockdown. Each dot represents the average value for one expression site. Bin size 20 kb. **d,** Virtual 4C analysis, viewpoint: SL-RNA locus (chr. 9). Relative interaction frequencies between the viewpoint and VSG expression sites is shown. Bin size 20 kb. **e,** Virtual 4C analysis, viewpoint: EP1 (chr. 10) or GPEET gene array (chr. 6). Relative interaction frequencies between the viewpoint and the SL-RNA locus are plotted. Bin size 20 kb. The analyses in **c** to **e** are based on Hi-C experiments with cells before and 24 h after VEX2 knockdown (n=3, the average is shown). **f,** Schematic model for monogenic VSG expression. A strong inter-chromosomal interaction between the *SL*-array and the active VSG gene facilitates spatial integration of transcription and mRNA maturation. VEX1 and VEX2 are primarily *SL*- and active-VSG associated, respectively, and sustain monogenic VSG expression by excluding other VSGs. The VSG-SL organelle is reconfigured upon activation of a different VSG.

To explore the role of VEX2 in controlling interactions between antigencoding genes and SL-RNA loci, we performed Hi-C analyses in VEX2-depleted cells. After 24 hours of VEX2 depletion, all previously silent expression sites displayed increased interaction frequencies with the SL-RNA locus. (**Fig. 4c**). The interaction between VSG expression site 3 and the SL-RNA locus showed the strongest increase. Notably, this is the expression site containing the most de-repressed VSG (*VSG-6*) following VEX2 depletion ^16^. Interaction frequencies of the active VSG expression site 1 with the SL-RNA locus remained unchanged, correlating with sustained and dominant *VSG-2* expression. Overall, SL-RNA interactions correlate with VSG transcript levels before and after VEX2 knockdown (**Extended Data Fig. 5a**).

Besides the VSG genes located in previously silent expression sites, expression site associated genes were also strongly upregulated following VEX2 knockdown ^16^. In line with this finding, we observed the largest increase in SL-RNA interactions for the regions upstream of the VSG gene in each de-repressed expression site, where expression site associated genes are located (**Fig. 4d, Extended Data Fig. 5b**). Thus, VEX2 restricts interactions between silent VSG expression sites, the expression site associated genes in particular, and the SL-RNA locus.

As a third group of RNA Pol l transcribed genes, insect stage specific procyclin genes are upregulated upon VEX2 depletion ^16^. Correlating with these data, following VEX2-knockdown, we found GPEET and EP1 procyclin genes to exhibit strongly increased interaction frequencies with the SL-RNA array and also with VSG expression sites (**Fig. 4e, Extended Data Fig. 5c**). Thus, our data suggest that VEX2 may have a dual function: specifically enhancing mRNA splicing of the VSG gene that is connected to the SL-RNA transcription compartment and, at the same time, excluding all other VSG expression sites and procyclin genes from the SL-RNA compartment to ensure monogenic VSG expression.

By combining proximity ligation and super-resolution microscopy, we were able to demonstrate spatial integration of the active VSG expression site and a genomic locus important for RNA maturation. Our data show that this supramolecular assembly is composed of a VSG transcription compartment with the active VSG gene, RNA Pol l and VEX2 and an SL-RNA transcription compartment with the SL-RNA array, RNA Pol II, VEX1 and the tSNAP complex, presumably together with other factors important for mRNA *trans*-splicing ^18^. Based on these findings we propose a model in which VSG choice is intimately associated with an inter-chromosomal interaction, bringing together two nuclear compartments to ensure efficient VSG mRNA processing at only one expression site (**Fig. 4f**). In the VSG transcription compartment, the VSG gene is transcribed by highly processive RNA Pol I, generating large amounts of VSG pre-mRNA that requires efficient processing to prevent premature degradation. In the SL-RNA transcription compartment, SL-RNA, an essential substrate for maturation of every mRNA, is produced. The close spatial proximity of the two compartments in a single locus provides a sufficiently high concentration of *trans*-splicing substrate to ensure the efficient maturation of highly abundant VSG transcripts. According to this model, VEX2 is the key molecule bridging the two compartments and excluding all but one VSG expression site from the SL-RNA transcription compartment, thereby ensuring expression of a single VSG gene per cell.

RNA processing as the limiting factor in monogenic VSG expression has been proposed previously, based on the observations that all VSG expression site promoters are active, yet processive polycistronic transcription and mature transcripts are specific to the single active expression site ^6,7^. Given the observation that co-transcriptional RNA processing can affect elongation rates in other organism ^21^, it will be interesting to determine whether efficient VSG mRNA processing will exert a positive feedback on transcription elongation along expression sites. Also, it was shown recently that splicing can activate transcription ^22^. It remains to be shown if factors located in the SL-RNA transcription compartment recruit the transcription machinery to the interacting VSG expression site and thereby enhance transcription. Notably, we also find other highly expressed housekeeping genes, such as core histones or tubulin, to associate with the SL-RNA locus (**Extended Data Fig. 6a**) and with VEX1 (**Extended Data Fig. 6b**). Thus, interactions with the SL-RNA locus may play a broader role for the regulation of gene expression in *T. brucei*.

While the importance of intra-chromosomal interactions in regulating gene expression has been shown in many complex eukaryotes, the significance of inter-chromosomal interactions has been questioned. Interactions between different chromosomes were thought to require complicated, possibly error-prone mechanisms to be re-established following mitosis. Yet, our data demonstrate a stable interaction between the active VSG gene, located either on chr. 6 or on an intermediate-sized chromosome, and the SL-RNA array locus on chr. 9. The interaction is also stably propagated during cell-division; despite being resolved during S phase, the interaction is re-established after replication. To our knowledge, these findings represent the first report of a selective inter-chromosomal interaction that connects transcription and mRNA splicing.

Protein condensates have recently emerged as important features that compartmentalize nuclear functions; transcription control by RNA polymerases, for example ^23^. Regulated switching between adjacent transcriptional and splicing condensates has been described in mammalian cells ^24^ and RNAs are major actors in facilitating genomic interactions and phase transitions ^25^. Given the recent finding that members of a family of helicases are global regulators of RNA-containing, phase-separated organelles ^26^, it is tempting to speculate that maintenance of VSG transcription-maturation compartment connectivity is similarly regulated by the putative RNA helicase VEX2 ^16^. We show here that in *T. brucei*, the assembly of two membrane-less nuclear condensates, each with a specific function in VSG gene transcription and RNA maturation, and both containing protein and RNA molecules ^12,18^, regulates monogenic VSG expression. By shaping a highly selective and specific genome architecture, VEX2 allows only one VSG gene to productively interact with the mRNA splicing compartment.

## Acknowledgements

We thank the Dundee Imaging Facility and J. Rouse for access to the Zeiss 880 Airyscan and Leica Confocal SP8 Hyvolution microscope, respectively, and S. Alsford (London School of Hygiene & Tropical Medicine) for the SNAP42 tagging construct. We further thank R. Cosentino and all members of the Siegel, Ladurner, Meissner and Boshart labs for valuable discussion, T. Straub (Core facility Bioinformatics, BMC) for providing server space and help with the data analysis, the Core Unit Systems Medicine, University of Würzburg for NGS.

## Author contributions

Experiments were designed by J.F., V.L., L.S.M.M., D.H. and T.N.S. and carried out by J.F., V.L. and L.S.M.M., unless indicated otherwise. Initial IFA experiments were performed by L.G.. RNAi, chemical inhibition and super resolution and other microscopy experiments were performed and analysed by J.F.. Hi-C experiments were performed by V.L. and L.S.M.M. and computational analysis was carried out by B.G.B., V.L. and L.S.M.M.. RNA-seq experiments were performed by V.L.; data analysis was carried out by V.L. and B.G.B.. ChIP-seq data analysis was carried out by S.H.. Funding was acquired by D.H. and T.N.S.. The work was supervised by D.H. and T.N.S.. The manuscript was written by J.F., V.L., D.H. and T.N.S. and edited by all other co-authors.

## Data and materials availability

Code and high-throughput sequencing data generated for this study have been deposited at GitHub (https://github.com/bgbrink/PRJEB35632) and in the European Nucleotide Archive (ENA) under primary accession number PRJEB35632, respectively. ChIP-seq data have also been deposited in the ENA (accession no. PRJEB25352). Processed data and results are available under https://doi.org/10.5281/zenodo.3628213.

## Methods

No statistical methods were used to predetermine sample size. The experiments were not randomized and investigators were not blinded to allocation during experiments and outcome assessment.

### *T. brucei* growth and manipulation

Bloodstream-form *T. brucei*, Lister 427 and 2T1 cells ^27^, both wild-type with respect to VEX1, VEX2 and SNAP42 subunits, were grown in HMI-11 medium and genetically manipulated using electroporation ^28^; cytomix was used for all transfections. Puromycin, phleomycin, hygromycin and blasticidin were used at 2, 2, 2.5 and 10 µg ml^-1^ for selection of recombinant clones; and at 1, 1, 1 and 2 µg ml^-1^ for maintaining those clones, respectively. RNAi experiments were undertaken through tetracycline induction at 1 µg ml^-1^. A double selection *T. brucei* cell line was used that derived from the Lister 427 bloodstream-form MITat 1.2 isolate ^19^. A neomycin resistance gene in VSG expression site 17 and a puromycin resistance gene in VSG expression site 1 allowed the selection for a homogenous cell population that either expressed *VSG-2* from expression site 1 or *VSG-13* from expression site 17. Cells were cultivated with either 10 µg ml^-1^ of neomycin (also referred to as N50 cells) or 0.1 µg ml^-1^ of puromycin (referred to as P10 cells). Established procyclic-form *T. brucei*, Lister 427 cells were grown in SDM-79 at 27 °C and genetically manipulated using electroporation as above. Blasticidin or hygromycin were used at 10 or 50 and 2 or 1 μg ml^-1^ for selection and maintenance, respectively.

### Plasmids

The VEX1 (Tb927.11.16920, 574 bp) ^15^, VEX2 (Tb927.11.13380, 471 bp) ^16^ and VEX1/VEX2 ^16^ RNAi cassettes were excised prior to electroporation by digesting with *Asc*I. The VEX1^12myc 15^ and SNAP42^12myc 29^ *C*-terminal tagging vectors were linearised with *Sph*I. The ^6myc^VEX2 and ^GFP^VEX2 *N*-terminal tagging vectors were linearised with *Xho*I. The SNAP42 GFP *C*-terminal tagging vector was made by replacing the 12 x c-myc tag and as also linearised with *Sph*I, respectively. Linearised RNAi constructs, under the control of tetracycline-inducible promoters, were transfected into 2T1 cells, which allow for targeting to a single genomic locus validated for robust inducible expression ^27^.

### ChIP-seq

ChIP-Seq was carried out as described in ^16^. Reads were aligned to the 11 curated megabase chromosomes from the TREU927 strain genome sequence ^30^, and a non-redundant set of BES and mVSG contigs from the Lister 427 strain ^31–33^ using bowtie2 ^34^ in very-sensitive alignment mode, and alignments were compressed and sorted using samtools ^35^. Bowtie2 attempted to align 54.0 and 49.9 million read pairs with 70.84 and 82.43 % success rates, respectively, resulting in 38.3 and 41.1 million aligned read pairs. PCR duplicate reads were removed using Picard MarkDuplicates (http://broadinstitute.github.io/picard/) resulting in 26.8 and 41.3 million aligned read pairs for analysis. Alignments were visually inspected with the Artemis genome browser ^36^. Circular plots (**Extended Data Fig. 3c**) were generated using the R library circlize ^37^ and bedgraph files for log2 fold change (**Fig. 3c, Extended Data Fig. 3c and Extended Data Fig. 6b**) were generated using deeptools2 ^38^. Bedgraphs were generated with 1kb bins and the option smoothLength 5000. Spliced leader RNA sequences were annotated using the sequences: Promoter:CGTTTCTGGCACGACAGTAAAATATGGCAAGTGTCTCAAAACTGCCTGTACAGCTTATTTTTGGGACACACCCATGCTTTC…Transcript…AACTAACGCTATTATTAGAACAGTTTCTGTACTATATTGGTATGAGAAGCTCCCAGTAGCAGCTGGGCCAACACACGCATT TGTGCTGTTGGTTCCCGCCGCATACTGCGGGAATCTGGAAGGTGGGGTCGGATGACCTC and the ‘transcript’ features were plotted with TSS and TES denoting the 5’ and 3’ extremities. Fold enrichment traces covering the spliced leader locus were calculated directly using deeptools bamCompare. Heat maps (**Extended Data Fig. 3c and Extended Data Fig. 6b**) were generated using deeptools2 bamCompare, computeMatrix and plotHeatmap ^38^ and resulting vector graphics files were then assembled into figures using Adobe Illustrator. Genomic regions for tandem genes and arbitrarily selected genes genes with a paralog count of 0 were assembled in bed files, using annotated mRNA sequences from TriTrypDB v5.1 of the TREU927 genome sequence. All scripts necessary to reproduce the ChIP-Seq analyses have been deposited together with the results of those analyses under https://doi.org/10.5281/zenodo.3628213.

### Protein blotting

Protein samples were run according to standard protein separation procedures, using SDS-PAGE. However, for VEX2 detection, the use of Bis-Tris gels with a neutral pH environment and a Bis-Tris/Bicine based transfer buffer (containing a reducing agent and 10% methanol) were critical for protein separation and transfer, respectively (NuPAGE, Invitrogen). Otherwise, western blotting was carried out according to standard protocols. The following primary antibodies were used: rabbit α-VEX2 (1:1,000), rabbit α-pol-I largest subunit ^15^ (1:500), rabbit α-VSG-2 (1:20,000), rabbit α-VSG-6 (1:20,000), mouse α-c-myc (Millipore, clone 4A6, 1:7,000), rabbit α-GFP (Abcam Ab290, 1:1,000) and mouse α-EF1α (Millipore, clone CBP-KK1, 1:20,000). We used horseradish peroxidase coupled secondary antibodies (α-mouse and α-rabbit, Biorad, 1:2,000). Blots were developed using an enhanced chemiluminescence kit (Amersham) according to the manufacturer’s instructions. Densitometry was performed using Fiji v. 2.0.0.

### Microscopy

Immunofluorescence microscopy was carried out according to standard protocols. For wide field microscopy (**Extended Data Fig. 4a**), the cells were attached to 12-well 5 mm slides (Thermo Scientific). For super resolution microscopy, the cells were attached to poly-L-lysine treated coverslips (thickness 1^1/2^ mm), stained and only then mounted onto glass slides. For colocalisation studies with Pol I we used antigen-retrieval. Prior to permeabilization, fixed cells were rehydrated in PBS for 5 min at RT, held at 95 °C for 60 s in freshly prepared antigen retrieval buffer (100 mM Tris, 5% urea, pH 9.5) and then washed 3 x 5 min in PBS at RT. Cells were mounted in Vectashield with DAPI (wide field) or stained with 1 µg ml^-1^ DAPI for 10 min and then mounted in Vectashield without DAPI (super resolution). In *T. brucei*, DAPI-stained nuclear and mitochondrial DNA were used as cytological markers for cell-cycle stage; one nucleus and one kinetoplast (1N:1K) indicates G1, one nucleus and an elongated kinetoplast (1N:eK) indicates S phase, one nucleus and two kinetoplasts (1N:2K) indicates G2/M and two nuclei and two kinetoplasts (2N:2K) indicates post-mitosis ^39,40^. Primary antisera were rat α-VSG-2 (1:10,000), rabbit α-VSG-6 (1:10,000), rabbit α-GFP (Invitrogen, 1:250; Abcam, 1:500), mouse α-myc (New England Biolabs, clone 9B11, 1:2,000), rabbit α-pol-I largest subunit ^15^ (1:100) or rabbit α-NOG1 ^41^ (1:500). The secondary antibodies were Alexa Fluor conjugated goat antibodies (Thermo Scientific): α-mouse, α-rat and α-rabbit, AlexaFluor 488 or Alexa Fluor 568 (1:1,000 or 1:2,000, for super resolution or wide field field microscopy, respectively). For wide field microscopy, cells were analysed using a Zeiss Axiovert 200M microscope with an AxioCam MRm camera and the ZEN Pro software (Carl Zeiss, Germany). The images were acquired as z-stacks (0.1-0.2 µm) and further deconvolved using the fast iterative algorithm in Zen Pro. For super resolution microscopy, cells were analysed using a Leica TCS SP8 confocal laser scanning microscope in Hyvolution Mode and the Leica Application Suite X (LASX) software (Leica, Germany) or a Zeiss 880 Airyscan and the Zeiss ZEN software (Carl Zeiss, Germany). The Hyvolution mode allows super resolution level images, general used settings: highest resolution / lowest speed; pinhole 0.5. All the super resolution images correspond to maximal 3D projections by brightest intensity of stacks of approximately 30 slices of 0.1 µm - Images with DNA in grey. For all quantifications, images were acquired with the same settings and equally processed. All the images were processed and scored using Fiji v. 2.0.0. ^42^. Sinefungin was applied at 2 µg ml^-1^ for 30 minutes at 37°C. VEX1, VEX2 and tSNAP foci and Pol I nucleolar and ESB signals could be detected in over 85-90% of nuclei. For all experiments involving RNAi, the knockdown was always verified by Western-Blot and the VSG dereppression phenotype confirmed by IFA and/or FACS analysis.

Counts in total cells or specific cell cycle phases were performed in >200 or >100 nuclei, respectively. All quantifications are averages or representative of at least two biological replicates and independent experiments. For the ESB / tSNAP localisation following VEX2 or VEX1 / VEX2 RNAi in **Fig. 4a-b** **and Extended Data Fig. 4c-d**, all the imaging and analysis was performed at 12 h post induction, a timepoint where there was sufficient VEX2 knockdown (**Extended Data Fig. 4b**) but both nucleolar Pol I and the ESB could be detected in > 85% of cells; the ESB is not detectable at later time points ^16^. Regarding the distance measurements between the ESB and tSNAP compartment (**Fig. 4c**), a control measurement (**Extended Data Fig. 4d**) was performed to make sure that the increase in the distance between the two protein condensates following VEX2 or VEX1 / VEX2 RNAi was not a mere consequence of a decrease in the ESB focus diameter. Moreover, the ESB / tSNAP localisation analyses following VEX RNAi or sinefungin treatment were restricted to G1 cells to exclude any cell cycle bias, as these protein condensates can separate during S phase (**Fig. 1c**).

### RNA-Seq

<u>RNA isolation:</u> The RNA-seq experiment and data analysis was performed as described previously ^43^, using three replicates each for *VSG-2* and *VSG-13* expressing cells. 45 million cells were harvested per replicate at 1,500 x g and 4 °C for 10 min. Cells were washed with 1× TDB (5 mM KCl, 80 mM NaCl, 1 mM MgSO_4_, 20 mM Na_2_HPO_4_, 2 mM NaH_2_PO_4_, 20 mM glucose pH 7.4). RNA isolation was performed using the NucleoSpin RNA kit (Macherey-Nagel; cat. no. 740955.10) according to the manufactureŕs instructions with minor changes. 3.8 µl of 1 M RNAse-free dithiothreitol (Sigma-Aldrich; cat. no. 10197777001) and 1 μl of 1:10 Ambion ERCC RNA Spike-In Mix (ThermoFisherScientific; cat. no. 4456739) was added to the cell lysis buffer prior to use. <u>Removal of ribosomal RNA:</u> rRNA was removed by hybridization as described previously ^43^. All solutions were kept free from nucleases. For each hybridization reaction, 2 μg of total RNA was mixed with 10 μl of formamide (SigmaAldrich; cat. no. F9037-100ML), 2.5 μl of 20× SSC (3 M NaCl, 0.3 M sodium citrate, the pH was adjusted to 7.0 with HCl), 5 μl of 0.005 M EDTA pH 8 (stock solution 0.5 M; ThermoFisherScientific; cat. no. AM9260G), 2.48 μl of 100 μM rRNA depletion mix (total 4 μg of oligos) and RNAse-free water (ThermoFisherScientific; cat. no. AM9938) to a total volume of 50 µl. Hybridization was performed for 5 min at 80 °C, ramp down to 25 °C at intervals of 5 °C per minute. Subsequently, 2 μl of RNAse-OUT (ThermoFisherScientific; cat. no. 10777019) and 50 μl of 1x SCC containing 20% formamide were added. Dynabead MyOne Streptavidin C1 beads (ThermoFisherScientific; cat. no. 65001) were prepared as recommended by the manufacturer for RNA applications and immobilization of nucleic acids. Three rounds of oligo capture were performed, using 120 μl (1.2 mg) of magnetics beads per round. The resulting supernatant, containing rRNA-depleted RNA, was purified using RNeasy MinElute CleanUp Kit (QIAGEN; cat. no. 74204). Depletion of rRNAs was evaluated on a 1.2% TBE-agarose gel. <u>cDNA synthesis, library preparation and sequencing:</u> Synthesis of cDNA was performed using NEBNext Ultra Directional RNA Library Prep Kit from Illumina (New England Biolabs; cat. no. E7420) according to the manufacturer’s instruction. The concentration of cDNA was measured using Qubit dsDNA HS Assay Kit (Invitrogen, cat. no. Q32854) and a Qubit 2.0 Fluorometer (Invitrogen; cat. no. Q32866). To generate strand-specific RNA-seq libraries, uracil excision and removing of the second strand was performed prior to conversion of Y-shaped adapters. Therefore, 3 μl of USER enzyme (New England Biolabs; cat. no. M5505) were mixed with 16 μl of adapter-ligated DNA, 1 μl of TruSeq PCR primer cocktail (50 μM) and 20 μl of KAPA HiFi HotStart ReadyMix (KAPA Biosystems, cat. no. KK2601). USER digestion was performed at 37 °C for 15 min, followed by the published amplification protocol. Library concentrations were determined in duplicate using Qubit dsDNA HS Assay Kit (Invitrogen, cat. no. Q32854) and a Qubit 2.0 Fluorometer (Invitrogen, cat. no. Q32866) and quantified using the KAPA Library Quantification Kit (KAPA Biosystems, cat. no. KK4824) according to the manufacturer’s instruction. Strand-specific RNA-sequencing libraries were sequenced in paired-end mode on an Illumina NextSeq 500 sequencer (2 × 75 cycles). <u>Processing of sequencing data:</u> The sequencing datasets were mapped to the TbruceiLister427_2018 genome assembly (release 43, downloaded from TriTrypDB ^44^) using BWA-mem ^45^. The alignments were converted from SAM to BAM format, sorted, indexed and filtered by alignment quality (q>0) using SAMtools version 1.9 ^35^. To visualize read coverage, the number of reads was normalized per billion mapped reads and coverage files were generated in the wiggle format using COVERnant version 0.3.0 with the subcommand *ratio* ^46^.

### In situ Hi-C

In situ Hi-C was performed as previously described ^5^. 2 × 10^8^ cells were collected per replicate and resuspended in 40 ml of 1× trypanosome dilution buffer (1× TDB; 0.005 M KCl, 0.08 M NaCl, 0.001 M MgSO_4_ × 7H_2_O, 0.02 M Na_2_HPO_4_, 0.002 M NaH_2_PO_4_ × 2H_2_O, 0.02 M glucose) or 1× PBS (insect stage cells). Cells were fixed in the presence of 1% formaldehyde for 20 min at room temperature by addition of 4 ml of 11% formaldehyde solution (50 mM Hepes-KOH pH 7.5, 100 mM NaCl, 1 mM EDTA pH 8.0, 0.5 mM EGTA pH 8.0, 11% formaldehyde). The reaction was stopped by addition of 3 ml of 2 M glycine and incubation for 5 min at room temperature and 15 min on ice. Cells were washed twice in 1× TDB for bloodstream form cells or 1× PBS for insect stage cells, respectively, and the cell pellet was snap-frozen in liquid nitrogen. Cells were resuspended in 1 ml of permeabilization buffer (100 mM KCl, 10 mM Tris pH 8.0, 25 mM EDTA) supplemented with protease inhibitors (1.5 mM pepstatin A, 4.25 mM leupeptin, 1.06 mM PMSF, 1.06 mM TLCK) and digitonin (200 μM final concentration) and incubated for 5 min at room temperature. Cells were washed twice in 1× NEBuffer3.1 (NEB, B7003S) and resuspended in 342 μl of 1× NEBuffer3.1. After addition of 38 μl of 1% SDS, and an incubation at 65 °C for 10 min, SDS was quenched by addition of 43 μl of 10% Triton-X 100 (Sigma). Incubation was continued at room temperature for 15 min. Another 35 μl of water, 13 μl of 10× NEBuffer3.1 and 100 units of MboI (NEB, R0147M) were added and the chromatin was digested at 37 °C overnight while shaking. To inactivate MboI, the sample was incubated at 65 °C for 20 min. Restriction fragments were biotinylated by supplementing the reaction with 60 μl of fill-in mix (0.25 mM biotin-14-dATP (Life Technologies, 19524016), 0.25 mM dCTP, 0.25 mM dGTP, 0.25 mM dTTP (Fermentas), 40 U of DNA polymerase I, large (Klenow) fragment (NEB, M0210)) and incubation at 23 °C for 4 h. The end-repaired chromatin was transferred to 665 μl of ligation mix (1.8% Triton-X 100, 0.18 mg BSA, 1.8× T4 DNA Ligase Buffer (Invitrogen, 46300018) and 5 μl of T4 DNA ligase (invitrogen, 15224025) were added. The ligation was performed for 4 h at 16 °C with interval shake. Crosslinks were reversed by adding 50 μl of 10 mg/ml proteinase K (65 °C for 4 h) following addition of another 50 μl of 10mg/ml proteinase K, 80 μl of 5 M NaCl and 70 μl of 10% SDS (65 °C, overnight). DNA was precipitated with ethanol and resuspended in 257 μl of TLE (10 mM Tris-HCl, 0.1 mM EDTA, pH 8.0). SDS was added to a final concentration of 0.1% and the sample was split among two tubes for sonication (Covaris S220; microtubes, 175 W peak incident power, 10% duty factor, 200 cycles per burst, 240 s treatment). The samples were recombined and the volume was adjusted to 300 μl with TLE. Fragments between 100 and 400 bp in size were selected using Agencourt AMPure XP beads (Beckman Coulter), according to the manufacturer’s instructions. The DNA fragments were eluted off the beads in 55 μl of TLE. For end-repair and biotin removal from un-ligated ends, 70 μl of end-repair mix was added (1× Ligation buffer (NEB), 357 μM dNTPs, 25U T4 PNK (NEB, M0201), 7.5U T4 DNA polymerase I (NEB, M0203), 2.5U DNA polymerase I, large (Klenow) fragment (NEB, M0210)) and incubated for 30 min at 20 °C and 20 min at 75 °C. To inactivate the enzymes, EDTA was added to a final concentration of 10 mM. To isolate biotin-labelled ligation junctions, 50 μl of 10 mg/ml Dynabeads MyOne Streptavidin C1 (Life Technologies, 65001) were washed with 400 μl of 1× Tween washing buffer (TWB; 5 mM Tris-HCl pH 7.5, 0.5 mM EDTA, 1 M NaCl, 0.05% Tween-20), collected with a magnet, resuspended in 400 μl of 2× binding buffer (10 mM Tris-HCl pH 7.5, 1 mM EDTA, 2 M NaCl) and added to the sample suspended in 330 μl TLE. Biotinylated DNA was bound to the beads by incubating the sample for 15 min at room temperature with slow rotation. Subsequently, the DNA-bound beads were captured with a magnet, washed twice with 400 μl of 1× binding buffer, washed once in 100 μl of 1× TLE T4 ligase buffer and resuspeded in 41 μl of TLE. For polyadenylation, 5 μl of 10× NEBuffer2.1, 1 μl of 10 mM dATP and 3 μl of 5 U/μl of Klenow fragment (3′→ 5′ exo (-)) (NEB, M0212) were added and the sample was incubated for 30 min at 37 °C followed by a deactivation step at 65 °C for 20 min. Beads were collected with a magnet, washed once with 400 μl 1× Quick ligation buffer (NEB, M2200) and resuspended in 46.5 μl of 1× Quick ligation buffer (NEB, M2200). 2.5 μl of DNA Quick ligase (NEB, M2200) and 0.5 μl of 50 μM annealed TruSeq adapters were added and incubated for 1 h at room temperature. Beads were separated on a magnet, resuspended in 400 μl of 1× TWB and washed for 5 min at room temperature with rotation. Beads were washed on the magnet with 200 μl 1× binding buffer and 200 μl of 1× NEBuffer2.1 and resuspended in 20 μl of 1× NEBuffer2.1. The library was amplified in eight separate reactions of 50 μl. Per reaction, 1.5 μl of 25 μM TruSeq PCR primer cocktail (TruSeq PCR primer cocktail_F, 5’-AATGATACGGCGACCACCGAG-3’; TruSeq PCR primer cocktail_R; 5’-CAAGCAGAAGACGGCATACGAG-3’), 25 μl of 2× Kapa HiFi HotStart Ready Mix (Kapa Biosystems, KR0370) and 21.5 μl of water were added to 2 μl of library bound to the beads. Amplification was performed as follows: 3 min at 95 °C, 5 cycles of 20 s at 98 °C, 30 s at 63 °C and 30 s at 72 °C, 1 cycle of 1 min at 72 °C, hold at 4 °C. The PCR reactions were pooled and the beads were removed from the supernatant using a magnet. The library was purified by addition of 1.5 volumes of Agencourt AMPure XP beads (Beckman Coulter), according to the manufacturer’s instructions. The sample was eluted off the beads using 25 μl of 1× TLE buffer, transferred to a fresh tube and the concentration was determined using Qubit (Qubit dsDNA HS Assay Kit, Thermo Fisher) and qPCR (KAPA SYBR FAST qPCR Master Mix, Kapa Biosystems), according to the manufacturer’s instructions. Library size distributions were determined on a 5% polyacrylamide gel. Paired-end 75-bp sequencing was carried out using the Illumina NextSeq 500 system with mid or high output NextSeq 500/550 kits v.2.5 according to the manufacturer’s instructions.

### Mapping of Hi-C reads and generation of interaction matrices

Reads were mapped to a modified version of the TbruceiLister427_2018 genome assembly (downloaded from TriTrypDB, release 43) containing the following modifications. For all Hi-C experiments, we masked a newly discovered misassembly in bloodstream expression site 2 (BES2) with Ns. For Hi-C experiments in 2T1-control ^27^ and VEX2 knockdown cells, we added the transfected constructs as separate contigs to the genome. The construct sequences, as well as the modified genome have been deposited together with the results of the analyses under https://doi.org/10.5281/zenodo.3568483. Mapping, filtering, normalization and read counting were performed by the mHi-C pipeline as described in ^17^. We modified the pipeline to be compatible with the *T. brucei* genome assembly and also incorporated a merging step for the individual replicates after the removal of duplicate reads, but before data normalization (step 4) in order to avoid the introduction of any bias by the merge. We chose ICEing as the normalization method and finally filtered the mHi-C outcome by the posterior probability of 0.6 (i.e. reads are assigned to a bin with a likelihood of at least 60%). Downstream analyses such as normalizing for the different ploidy within the *T. brucei* genome assembly, have been implemented with in-house scripts. The digestion of the reference genome with the restriction site has been implemented using HiC-Pro Utilities ^47^. All scripts necessary to reproduce the Hi-C analyses can be found at: https://github.com/bgbrink/PRJEB35632.

### Virtual 4C analysis

To visualize interactions between one genomic region (viewpoint) and all other genomic sites, relevant bins were extracted from a 20-kb or 50-kb Hi-C matrix. An average interaction value for every genomic bin was calculated if the viewpoint regions spanned more than one bin. The coordinates that define the different viewpoints used in this study are shown in Data S1 sheet 2. To determine the relative interaction frequency of a viewpoint with chromosome cores and sub-telomeres, the average interaction frequency of the viewpoint with each chromosome core and sub-telomere was calculated based on the relative interaction frequencies extracted by virtual 4C analysis. The ratio between the average interaction frequency (core) and the average interaction frequency (sub-telomeres) was calculated for each chromosome and plotted as one dot. The virtual 4C analysis has been implemented using HiC sunt dracones (https://doi.org/10.5281/zenodo.3570496). All scripts necessary to reproduce the Hi-C analyses can be found at: https://github.com/bgbrink/PRJEB35632.

### Statistical analysis

All statistical analysis was performed using GraphPad Prism Software (version 7.0). A detailed summary of *n* and *p* values for all the analyses performed in this study is provided in Data S1 sheet 3.

### Resources and Reagents

All unique materials are available on request.

## Data S1. (separate file)

Sheet 1. ChIP-Seq data for VEX1^myc^ expressing bloodstream form *T. brucei*

Sheet 2. Coordinates of the viewpoints used in the virtual 4C analyses

Sheet 3. Summary of all statistical analyses

## Extended Data

**Extended Data Fig. 1.**
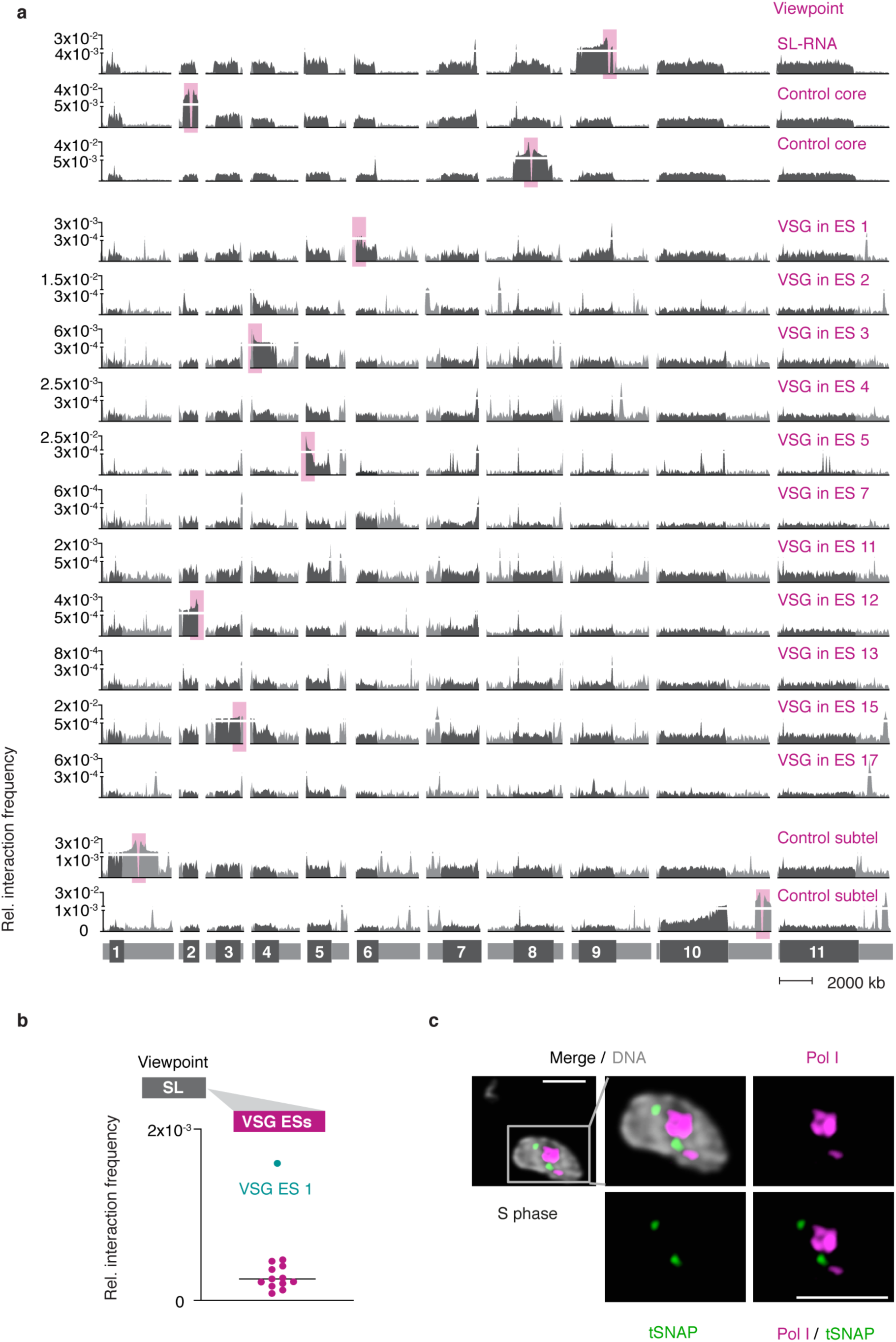
Genome-wide interaction frequencies of VSG genes in expression sites and the SL-RNA locus. **a,** Hi-C (virtual 4C) analysis with locations of viewpoints marked by pink boxes. Viewpoints VSG ES 4, 7, 11 and 17 are located on intermediate chromosomes that are not depicted in this figure. Interaction frequencies between each viewpoint and the 11 mega-base chromosomes are shown. Chromosome cores, dark grey; sub-telomeric regions, light grey. The hemizygous sub-telomeric regions of each chromosome are displayed in the following order: 5’(haplotype A)–5’(haplotype B)–diploid chromosome core–3’(haplotype A)–3’(haplotype B). Bin size 50 kb. Virtual 4C analyses in **a-b** are based on Hi-C experiments of *VSG-2* expressing cells (n=2, the average is shown). **b,** Virtual 4C analysis, viewpoint: SL-RNA locus (chr. 9). Relative interaction frequencies between the viewpoint and the active VSG ES 1 (cyan) and inactive VSG ESs (magenta) is shown. Each dot represents the average value for one expression site. Bin size 20 kb. **c,** Immunofluorescence-based colocalisation studies of tSNAP^myc^ (SL-RNA transcription compartment) and a nucleolar and active-*VSG* transcription compartment marker (Pol I, largest subunit) using super resolution microscopy. DNA was counter-stained with DAPI; the images correspond to maximal 3D projections of stacks of 0.1 μm slices and are representative of two biological replicates and three independent experiments; scale bars 2 μm.

**Extended Data Fig. 2.**
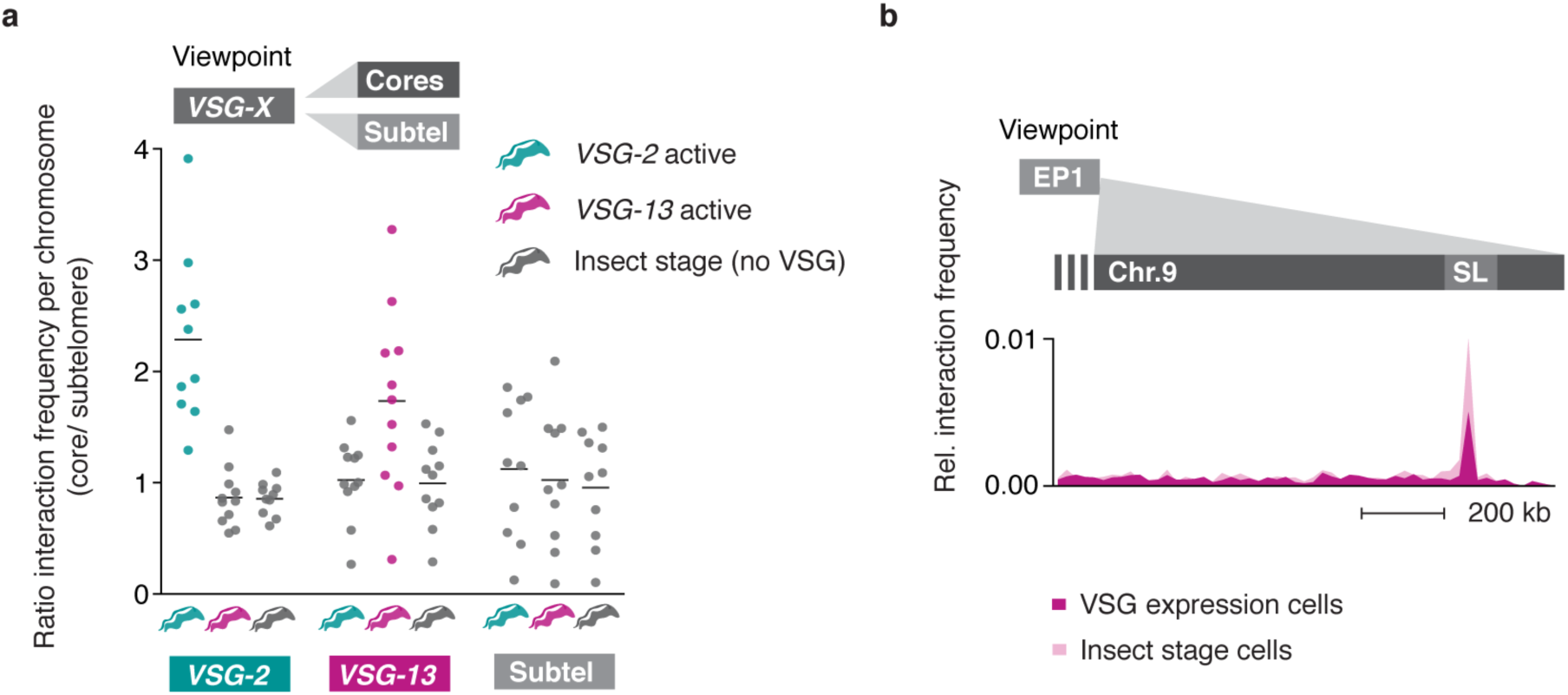
Changes in DNA-DNA interactions following a change in VSG isoform expression. **a,** Hi-C (virtual 4C) analysis, viewpoints *VSG-*2, *VSG-13* and a sub-telomeric control region. Each dot represents the ratio of: the average interaction frequency of the viewpoint with the chromosome core / the average interaction frequency with the sub-telomeres. One dot per chromosome is plotted. The black bar marks the median ratio per viewpoint. Bin size 50 kb. **b,** Virtual 4C analysis, viewpoint: EP1 gene array (chr. 10). Relative interaction frequencies between the EP1 array and the SL-RNA locus are plotted. Bin size 20 kb.

**Extended Data Fig. 3.**
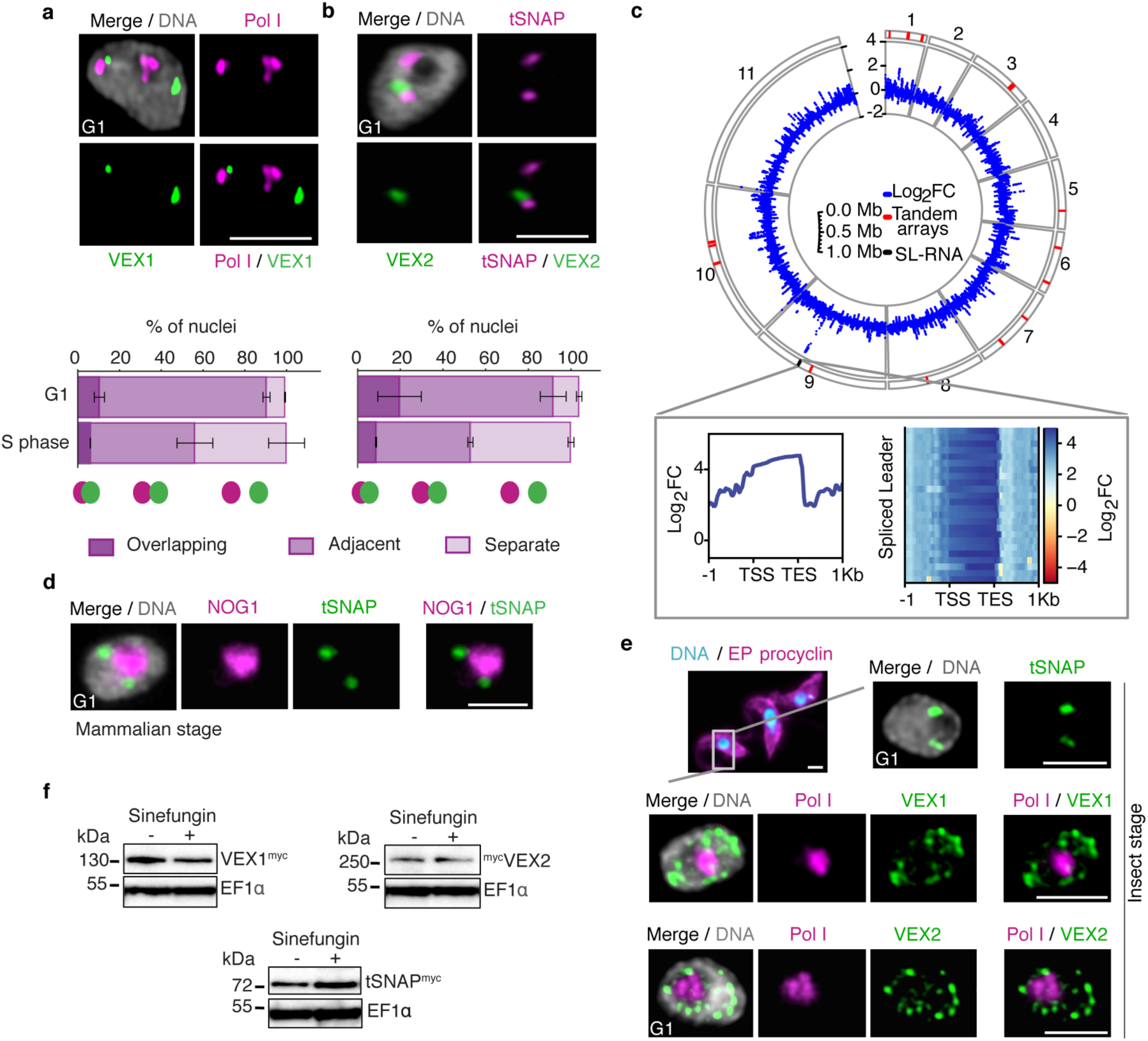
The VEX complex associates with both the active-VSG and the *Spliced Leader* (*SL*)-locus in a cell cycle and developmental stage-dependent manner. **a-b,** Immunofluorescence-based colocalisation studies of VEX1^myc^ / Pol I and ^GFP^VEX2 / tSNAP^myc^ in bloodstream form cells. tSNAP and Pol I were used as markers for the SL-RNA and VSG transcription compartments, respectively. The stacked bar graphs depict proportions of nuclei with overlapping, adjacent or separate signals and values are averages of two independent experiments (≥100 nuclei for G1 and S phase cells); detailed *n* and *p* values are provided in Data S1 sheet 3. **c,** VEX1^myc^ chromatin immunoprecipitation followed by next generation sequencing (ChIP-seq) analysis. The circle plot represents log2 fold change of ChIP versus Input of non-overlapping 1 kbp bins of the 11 megabase chromosomes; outside track shows tandem arrays (red) and the SL-RNA locus (black). An inset zooming on the SL-RNA locus is depicted: metagene plot (left-hand side) and heat-map (right-hand side) of SL-gene loci. Bin size 300 bp. **d,** Immunofluorescence-based colocalisation studies of tSNAP^myc^ and a nucleolar marker (NOG1) in bloodstream forms. **e,** Localisation of tSNAP^GFP^ and colocalisation studies of VEX1^myc^ or ^myc^VEX2 and Pol I in procyclic forms (insect-stage), using immunofluorescence. Procyclic forms do not express VSGs whereas procyclins are the major surface glycoprotein. Images in **a-b** / **d-e** were obtained with super resolution microscopy and correspond to maximal 3D projections of stacks of 0.1 μm slices; DNA was counter-stained with DAPI; scale bars 2 μm. **f,** Western-blot analysis of VEX1^myc^, ^myc^VEX2 and tSNAP^myc^ before and after sinefungin treatment (5 μg ml^-1^ for 30 min at 37°C), which blocks *trans*-splicing in trypanosomes. Data in **a-b** and **d-f** are representative of at least two independent biological replicates and two independent experiments.

**Extended Data Fig. 4.**
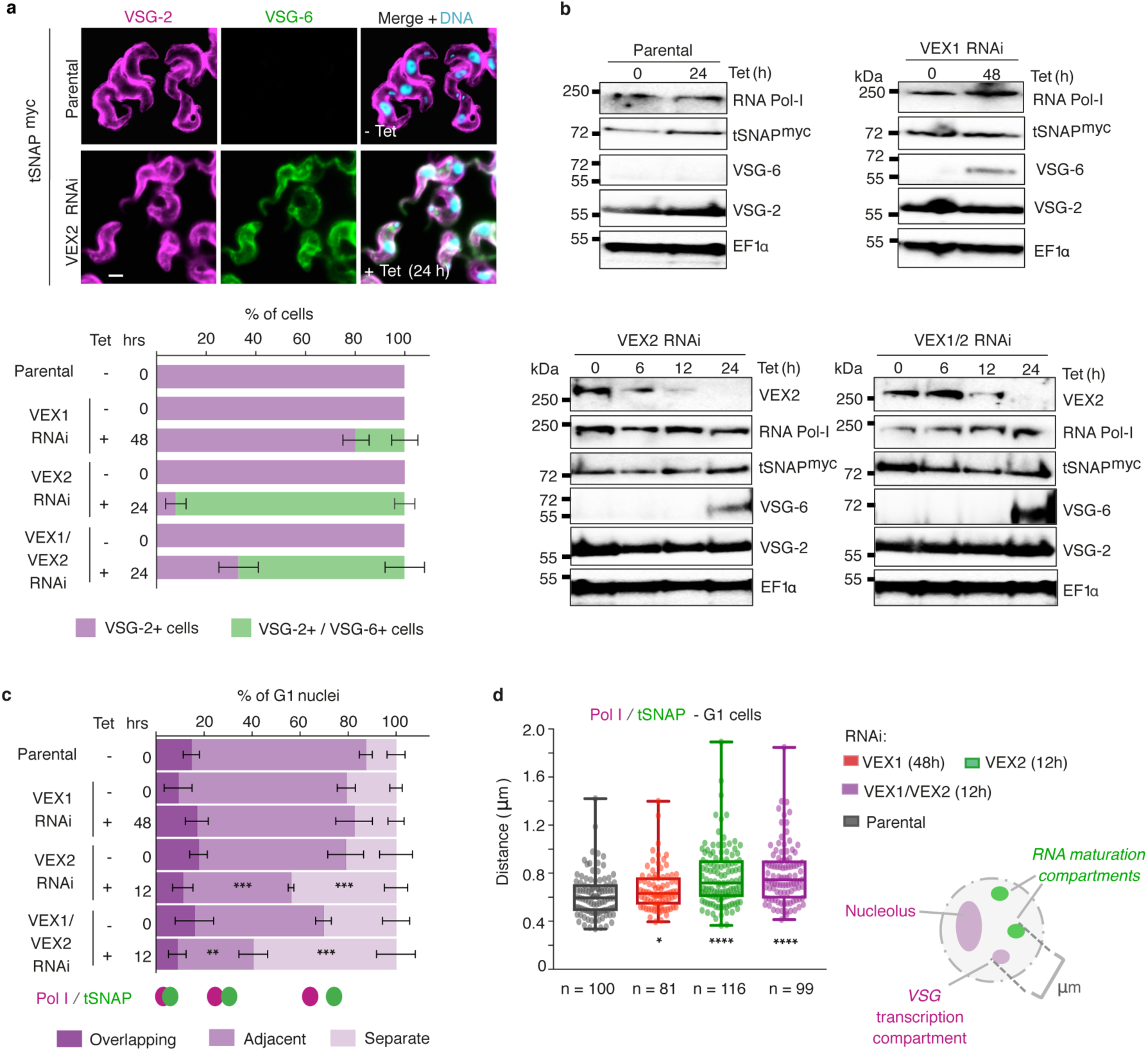
Pol I and tSNAP expression and localisation following knockdown of the VEX complex. **a,** Immunofluorescence-based analysis of VSG expression following tetracycline (Tet) inducible VEX1 knockdown, VEX2 knockdown or VEX1/VEX2 knockdown. VSG-2 (magenta) is the active-*VSG* and VSG-6 (green) is a silent-*VSG* in this strain. The stacked bar graph depicts percentages of VSG-2 single positive cells and VSG-2/VSG-6 double positive cells; values are averages of two independent experiments and two biological replicates. DNA was counter-stained with DAPI; scale bar 2 μm. **b,** Western-blot analysis of VEX2, Pol I, tSNAP^myc^, VSG-6 and VSG-2 expression following VEX1, VEX2 or VEX1/VEX2 knockdown. EF1α was used as a loading control. The data is representative of two independent experiments and two biological replicates. **c-d**, Immunofluorescence-based colocalisation studies of tSNAP^myc^ and a nucleolar and active-*VSG* marker (Pol I, large subunit). The stacked bar graph in **c** depicts proportions of G1 nuclei with tSNAP^myc^ / Pol I overlapping, adjacent or separate signals following tetracycline (Tet) inducible VEX1 (48 h), VEX2 (12 h) or VEX1/VEX2 knockdown (12 h). tSNAP^myc^ / active-*VSG* localisation were not monitored beyond 12 h following VEX2 and VEX1/2 knockdown as Pol I signal drops below detection at later time-points. The values are averages of two independent experiments and two biological replicates (≥100 G1 nuclei). In the box plot in **d**, the distance between the edges of the ESB and tSNAP foci was measured in > 81 G1 nuclei. The centre lines show the medians; box limits indicate the 25th and 75th percentiles; whiskers extend from maximal to the minimal values; all data points are shown. In **a/c**, error bars, SD. In **c-d**, knockdown conditions were compared to parental cells using two-tailed paired (**c**) or unpaired (**d**) Student’s *t*-tests: *, *p* < 0.05; **, *p* < 0.01; ***, *p* < 0.001; ****, *p* < 0.0001. Detailed *n* and *p* values are provided in Data S1 sheet 3.

**Extended Data Fig. 5.**
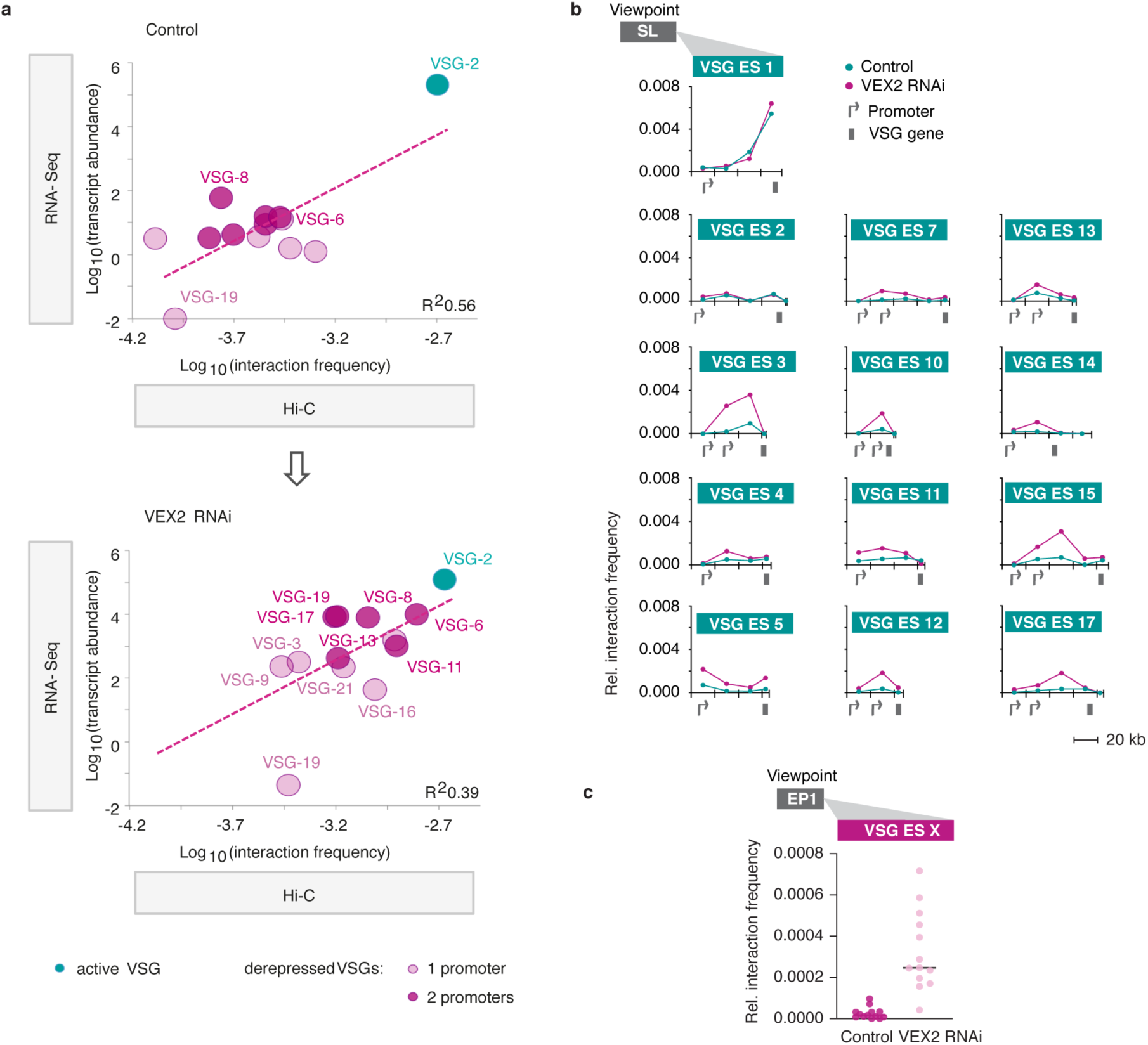
Genome-wide changes in VEX2 depleted cells. **a,** Correlation between the average interaction frequency of VSG expression-sites as viewpoint with the SL-RNA locus and VSG expression in reads per kilobase per million (RNA-seq data from ^16^) in control and VEX2-depleted cells. **b,** Hi-C (virtual 4C) analysis, viewpoint: SL-RNA locus (chr. 9). Relative interaction frequencies between the viewpoint and VSG expression sites are shown. Bin size 20 kb. **c,** Virtual 4C analysis, viewpoint: EP1 gene array (chr.10). Relative interaction frequencies between the viewpoint and the VSG expression sites are shown. Each dot represents the average value for one expression site. Bin size 20 kb.

**Extended Data Fig. 6.**
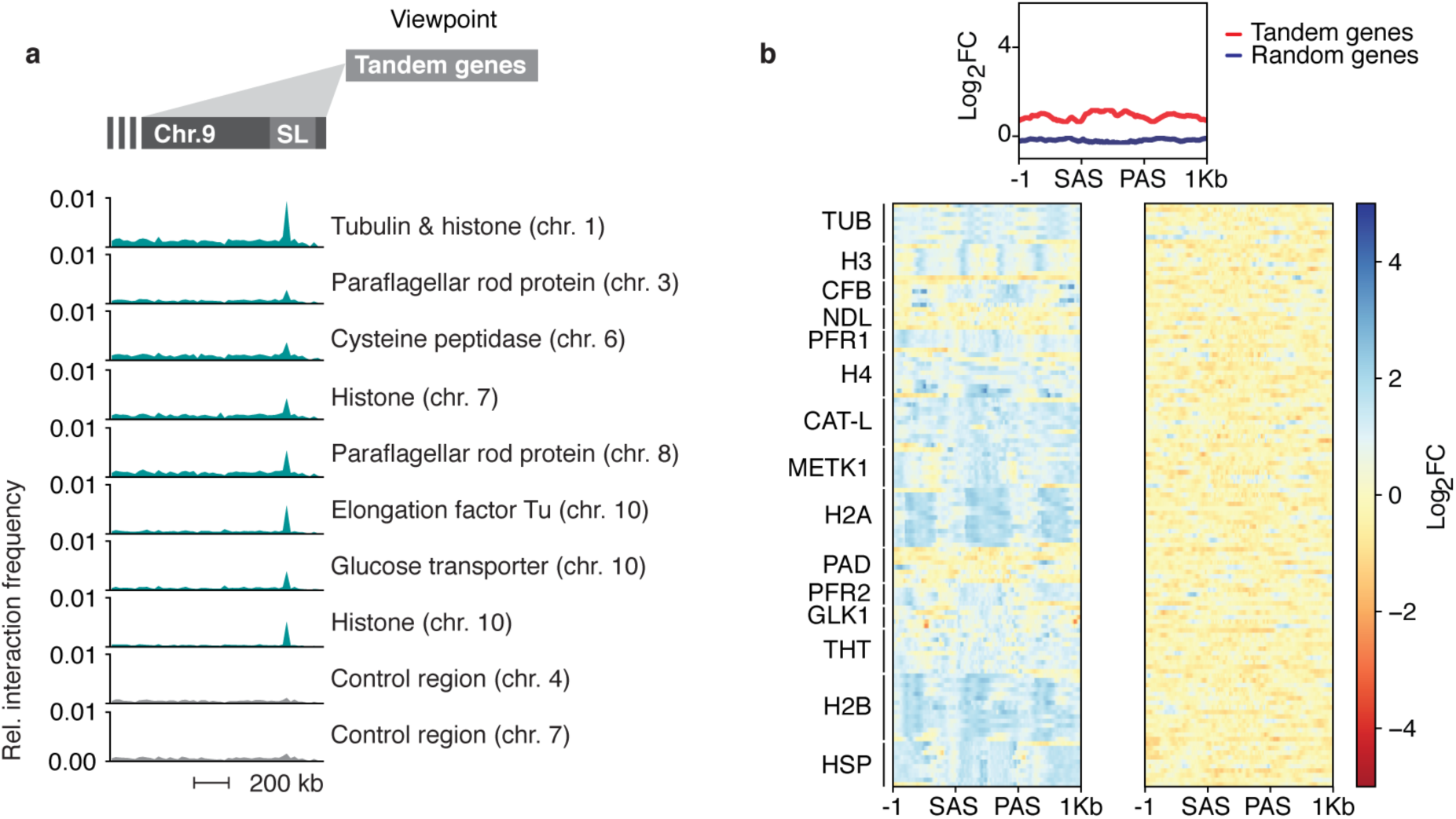
Tandem arrays interact at lower frequency with the *Spliced Leader* (*SL*)-locus. **a,** Hi-C (virtual 4C) analysis, viewpoint: different tandem gene arrays and control sites. Relative interaction frequencies between the different viewpoints and the SL-RNA locus are plotted. Bin size 20 kb. **b,** VEX1^myc^ chromatin immunoprecipitation followed by next generation sequencing (ChIP-seq) analysis. Top panel, metagene plot for tandem genes compared to randomly selected genes with no paralogues. Lower left, heat map of tandem genes. Lower right, randomly selected genes with no paralogues.

